# Influence of the culture media on the aspect and growth of oilseed rape fungal pathogens

**DOI:** 10.1101/2020.05.25.114314

**Authors:** Magali Ermel, Stéphane Jumel, Lydia Bousset

## Abstract

While culturing fungal pathogens on artificial media, the phenotype, i.e. colony aspect, growth kinetics and in some case the pigment production can be used as discrimination criteria. Our aim was to enable the comparison between the growth kinetics of 8 fungal species, cultured simultaneously on 3 different media, and imaged under standardised conditions. Using a camera stand, taking pictures was quick and easy, producing homogeneous pictures, facilitating their comparison. The colony growth kinetics varied widely across the 8 species. The mycelium aspect and pigment production depended on the media and changed over time. Having standardised the imaging setup and grown species simultaneously allows proposing reference sets of pictures.

## Introduction

While studying fungal pathogens, their culture on artificial media might be required, for example for diagnostic purpose, or to keep strains in collection. When fungi are isolated from natural populations or sub cultured from collections, species diagnostic is important to avoid the risk of co-infections by several species, and to discard contaminations.

While molecular markers or the observation of spore shapes under binocular lens allows accurate species diagnostic, simple visual control is done during isolation and sub culturing. The phenotype, i.e. colony aspect, growth kinetics and in some case pigment production can be used as discrimination criteria. However, the nutrient composition and physicochemical factors such as pH and oxydo-reduction status of the media can modify the colony aspect (Bousset et al. 2019). When the fungi do not have specific requirements for growth or sporulation, each research group uses different artificial media and get used to recognise species on this specific medium. Transposing this knowledge on alternative media is difficult.

Our aim was to enable the comparison between the growth kinetics of 8 fungal species, cultured simultaneously on 3 different media, and imaged under standardised conditions.

## Materials and Methods

### Artificial media

As none of the species had specific requirements, we compared the three media in use in the research groups working with oilseed rape fungal pathogens: Malt Agar, Potato Dextrose Agar (PDA) and V8 vegetable juice media.

Malt medium was prepared with 20g malt extract and 20g agar; PDA medium was prepared with 20g Potato Dextrose Broth and 20g agar; V8 medium was prepared with 20ml V8 vegetable juice and 20g agar. Each was completed to 1 litre with deionised water, then bottles were autoclaved for 20 mn at 120°C. Once cooled down below 70°C, 1 ml of Streptomycin solution 10% weight/volume was added (for a final concentration of 0,1 g.l^−1^) and 5 cm diameter Pétri dishes were poured.

### Studied species and growth conditions

We worked with 8 species of oilseed rape fungal pathogens: *Alternaria brassicae*, *Leptosphaeria biglobosa*, *L. maculans*, *Mycosphaerella brassicicola*, *Pseudocercosporella capsellae*, *Pyrenopeziza brassicae*, *Sclerotinia sclerotiorum* and *Verticillium longisporum*.

Strains issued from our collection were sub cultured on malt medium, then a 5 mm diameter plug was punched from the culture front and placed in the centre of each Pétri dish. Sub culturing was performed simultaneously on the 3 artificial media and for 7 of the species; the sub culturing of *L. maculans* was repeated independently. Dishes were grown in the laboratory, under neon lights of the room. Average temperature was 21,4°C ± 0,8 and relative moisture was 43,9 ± 3,8.

### Imaging the dishes

At several times during the colony growth, standardised pictures of the Pétri dishes were taken. Opening the dishes under the laminar flow allowed avoiding dew droplets on the dish cover, while preserving a sterile environment. Dish was placed on a white paper, with a ruler and with labels for species name and for time after colony transfer. A checker chart (white, grey and black pieces) with appropriate texture and colour for white balancing was included on the picture. Two workshop LED lamps provided light from both sides of the dish, places within the lower 45° angle to avoid reflection (Figure 1).

**Figure 1:**
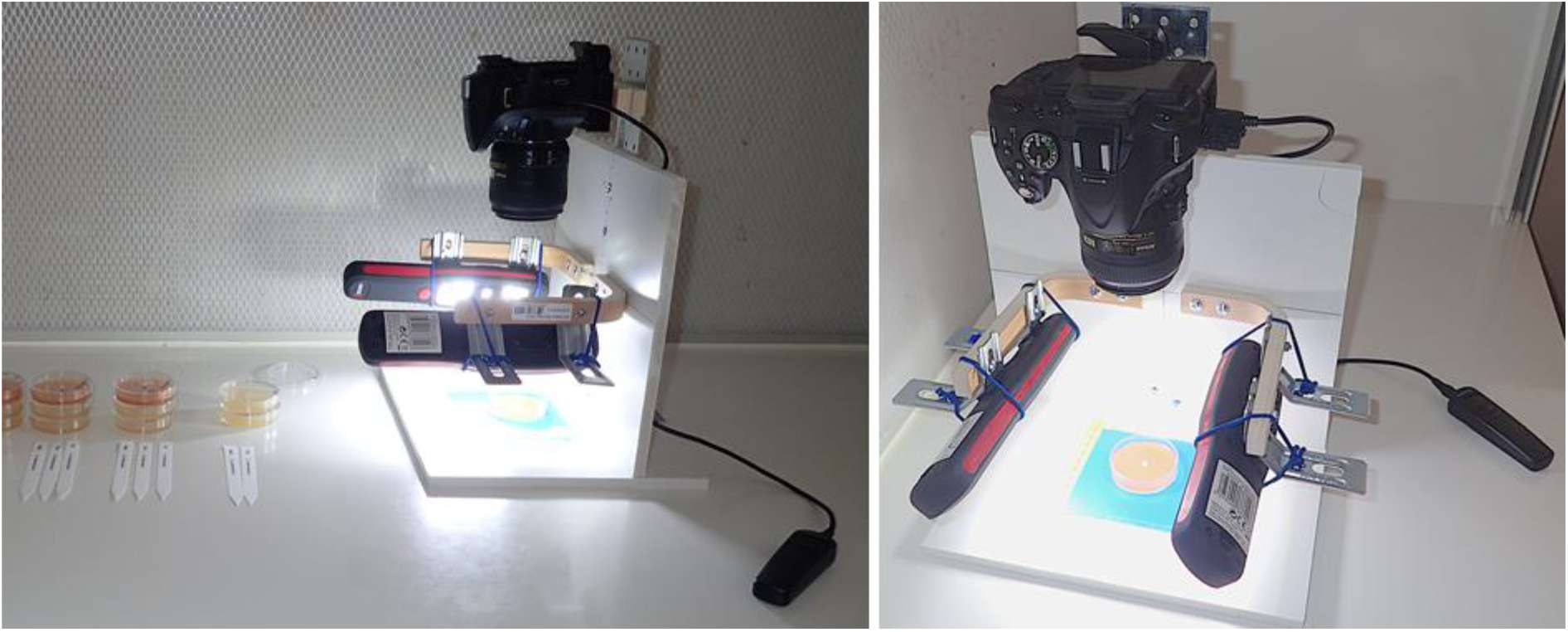
Camera setup under the laminar flow.

Pictures were taken with a Nikon D520 camera with a AF-S DX Micro Nikkor 40mm 1:2.8G macro lens, attached on a specially designed stand to allow the air flow, keeping the opened dish sterile. The distance between the front of the lens and the dish was 20 cm ad we used a remote control. Light measurement and focus were in the image centre, on the colony. Aperture was set to F22 for maximal depth of field, with 125 iso sensitivity and daylight white balance. Images were saved in RGB format 6000 × 4000 pixels.

### Image post-processing

As picture conditions were slightly different among the series, we adjusted the white balance and extracted the background on each image. This post-processing was done with 3 macro in the ImageJ software, successively applied to the folder containing the images. The first macro does the white balance adjustment using the checker chart present on each picture. The second macro crops pictures in batch mode, to a specified size. The third macro creates a mask for the Pétri dish then removes the background.

## Results

### Imaging

Using the camera stand with LED lamps made it possible to standardise the picture magnification, with homogeneous light (Figure 2a).

**Figure 2:**
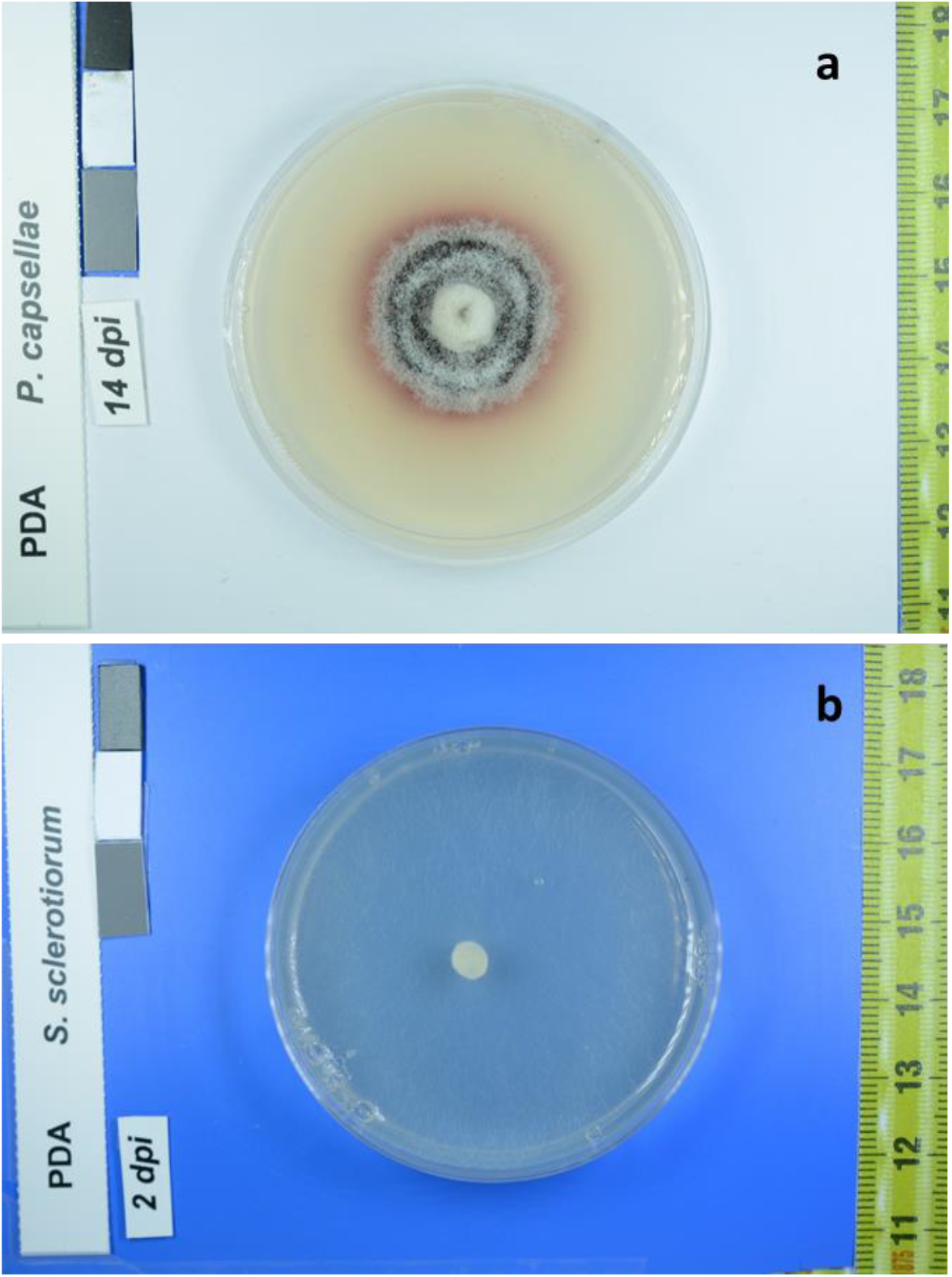
Examples of Pétri dish pictures with the ruler, the checker chart for white balancing and the identification labels. a) *Pseudocercosporella capsellae* on PDA, on white background 14 days after colony transfer; b) *S. sclerotiorum* on PDA, on blue background 2 days after colony transfer.

### Colony growth

Colony growth was very different depending on the species and it depended on the agar medium. Only the relative growth was compared across the 8 species studied.

*S. sclerotiorum* and *A. brasicae* had the fastest growth (Figure 3, 4). *S. sclerotiorum* filled the dish in 2 days. Its mycelium was difficult to see on the white background, before it changed to a cotton-like appearance around 4 dpi. The early growth was better visualised on a blue background (Figure 2b). *A. brassicae* spread over half of the dish diameter and its colony took a dark coloration in 4 days. It covered the whole dish in about a week.

**Figure 3:**
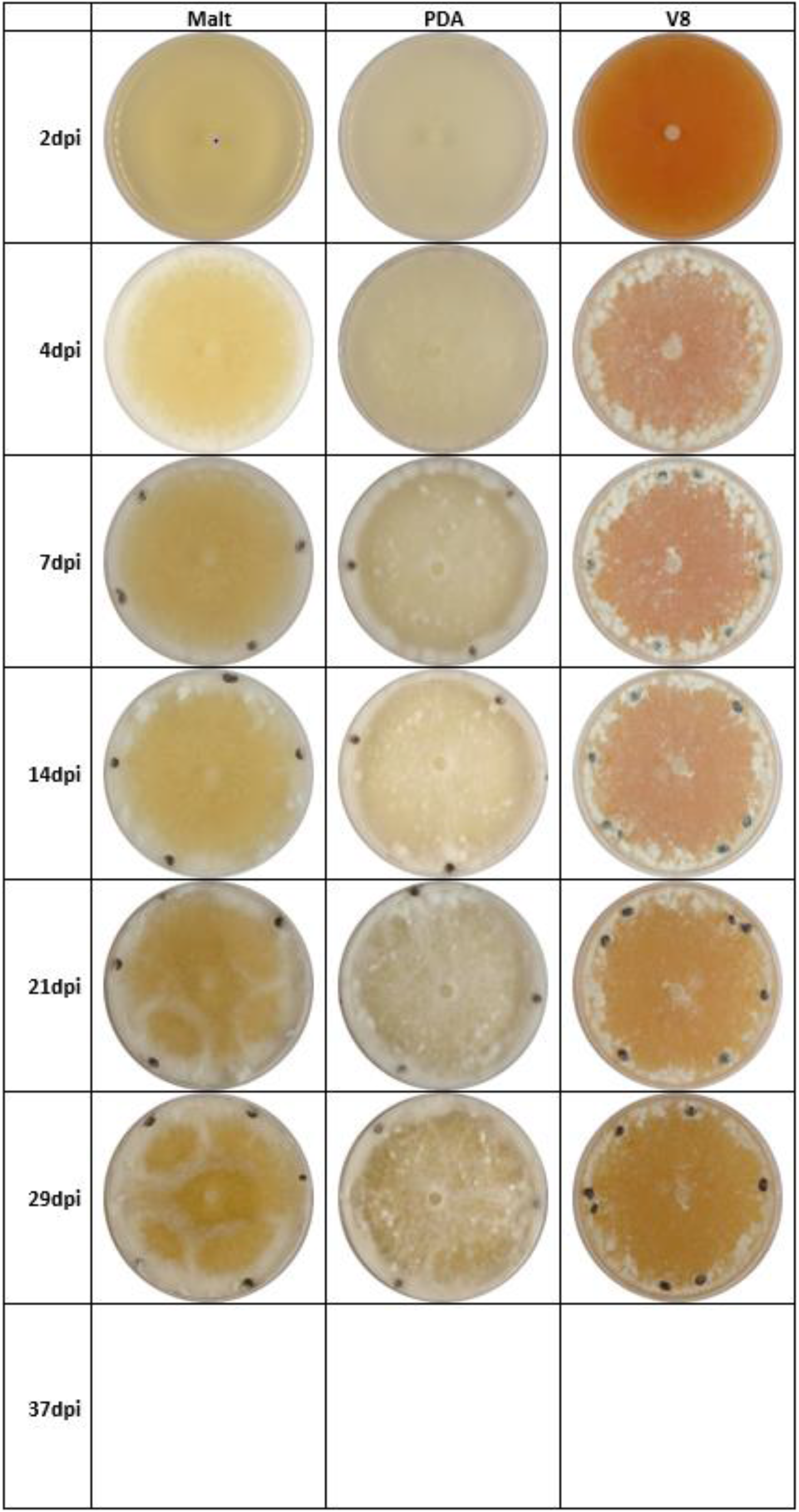
Growth kinetics of the same *Sclerotinia sclerotiorum* strain on malt, PDA and V8 agar media.

**Figure 4:**
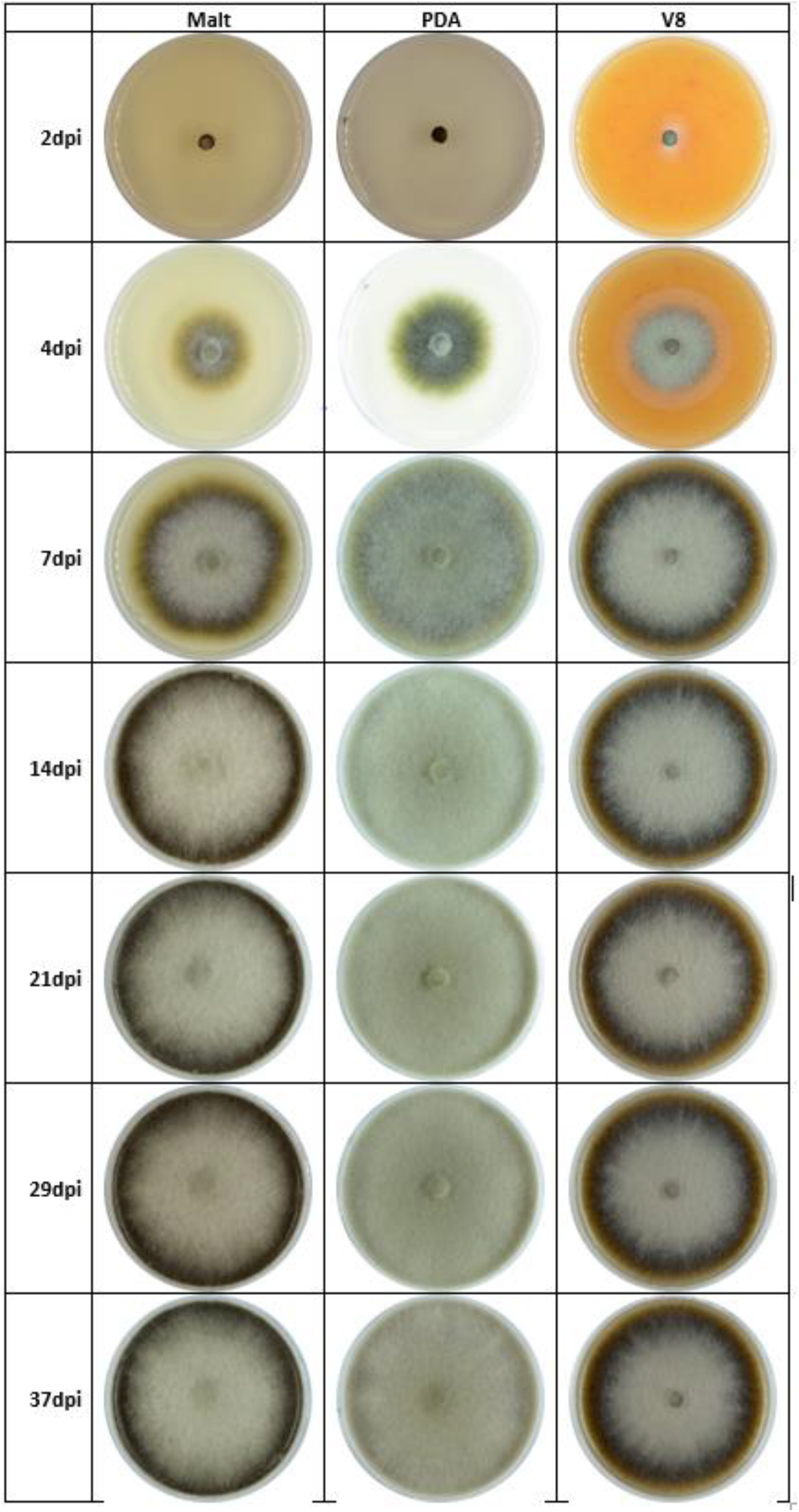
Growth kinetics of the same *Alternaria brassicae* strain on malt, PDA and V8 agar media.

*V. longisporum*, *L. biglobosa* and *L. maculans* had a fair growth after a week and their colonies acquired coloration during the second week of growth (Figure 5, 6, 7).

**Figure 5:**
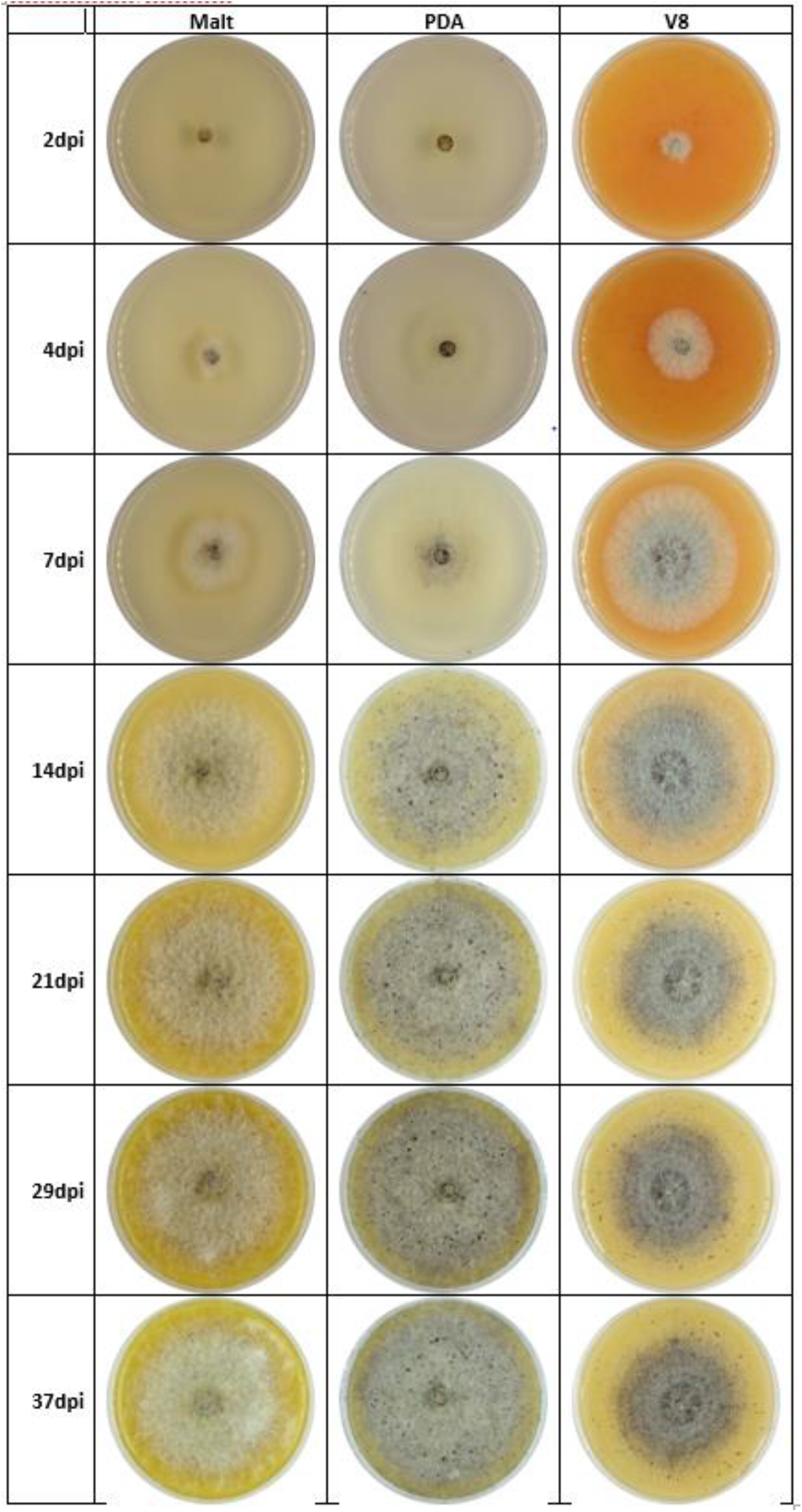
Growth kinetics of the same *Leptosphaeria biglobosa* strain on malt, PDA and V8 agar media.

**Figure 6:**
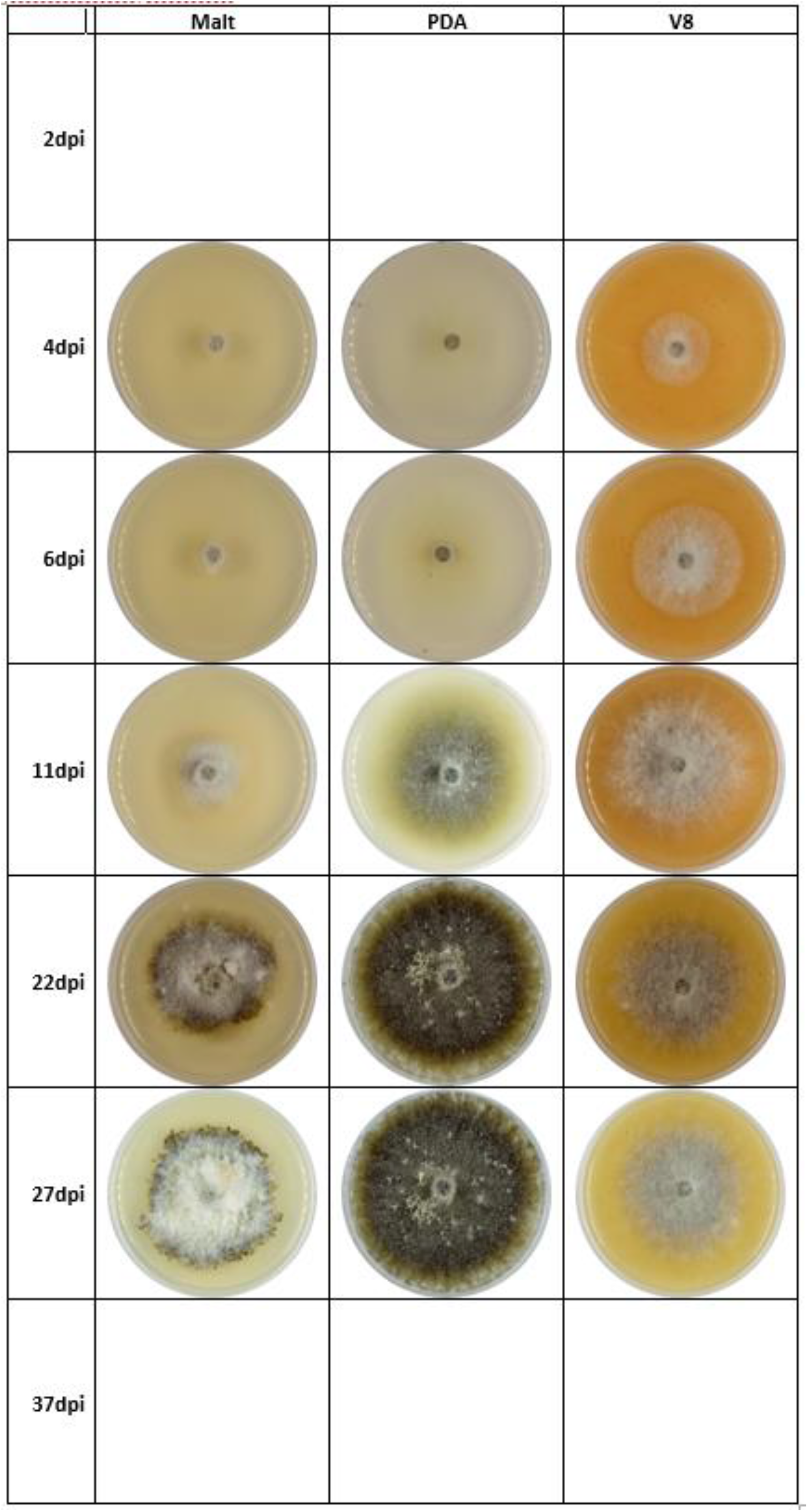
Growth kinetics of the same *Leptosphaeria maculans* strain on malt, PDA and V8 agar media. Note that pictures were taken at different times than the other species.

**Figure 7:**
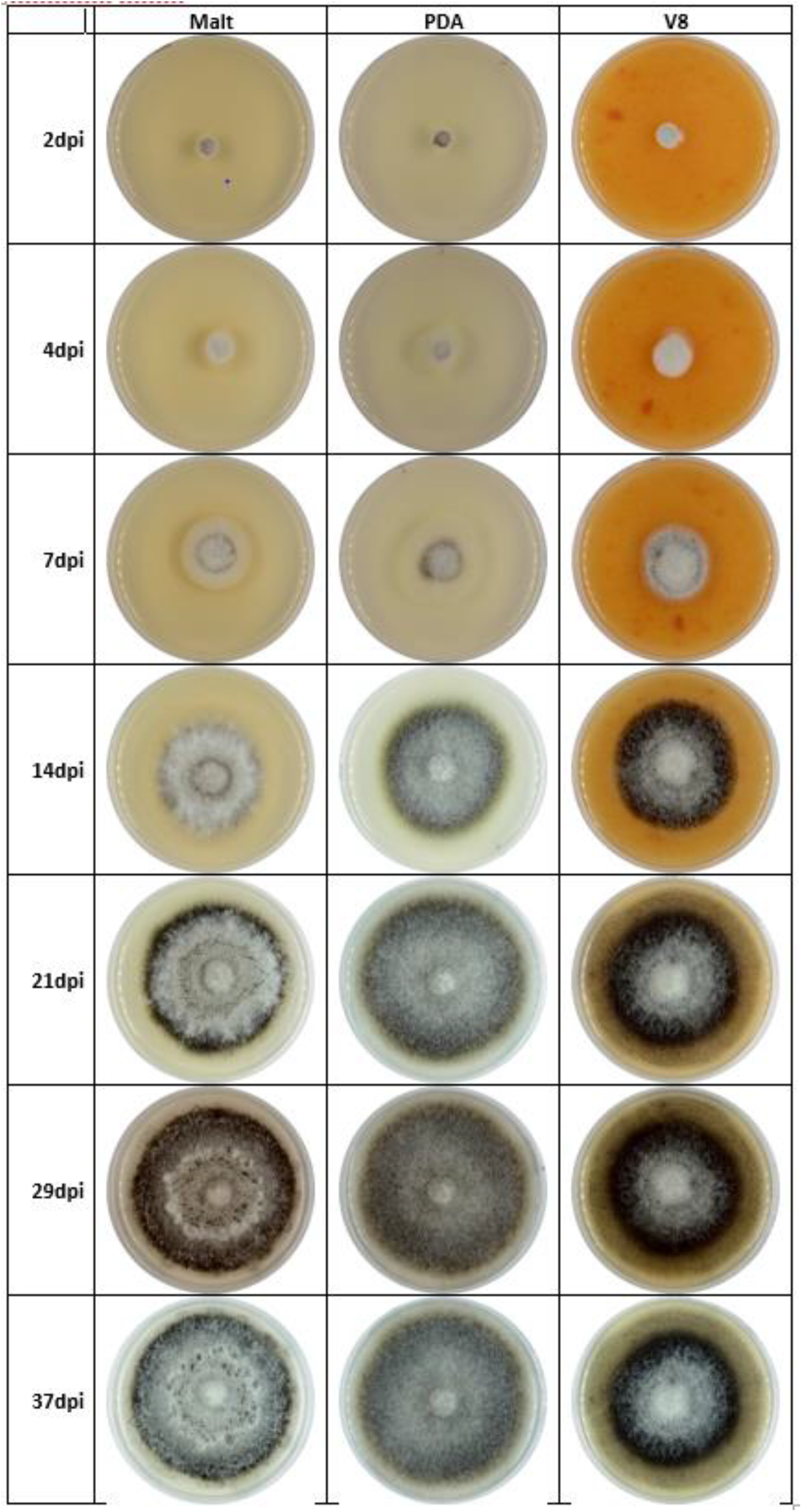
Growth kinetics of the same *Verticillium longisporum* strain on malt, PDA and V8 agar media.

*M. brassicae* and *P. capsellae* grew slower (Figure (8, 9). At 2 weeks, their colonies had only 1 to 2 cm diameter, depending on the medium.

**Figure 8:**
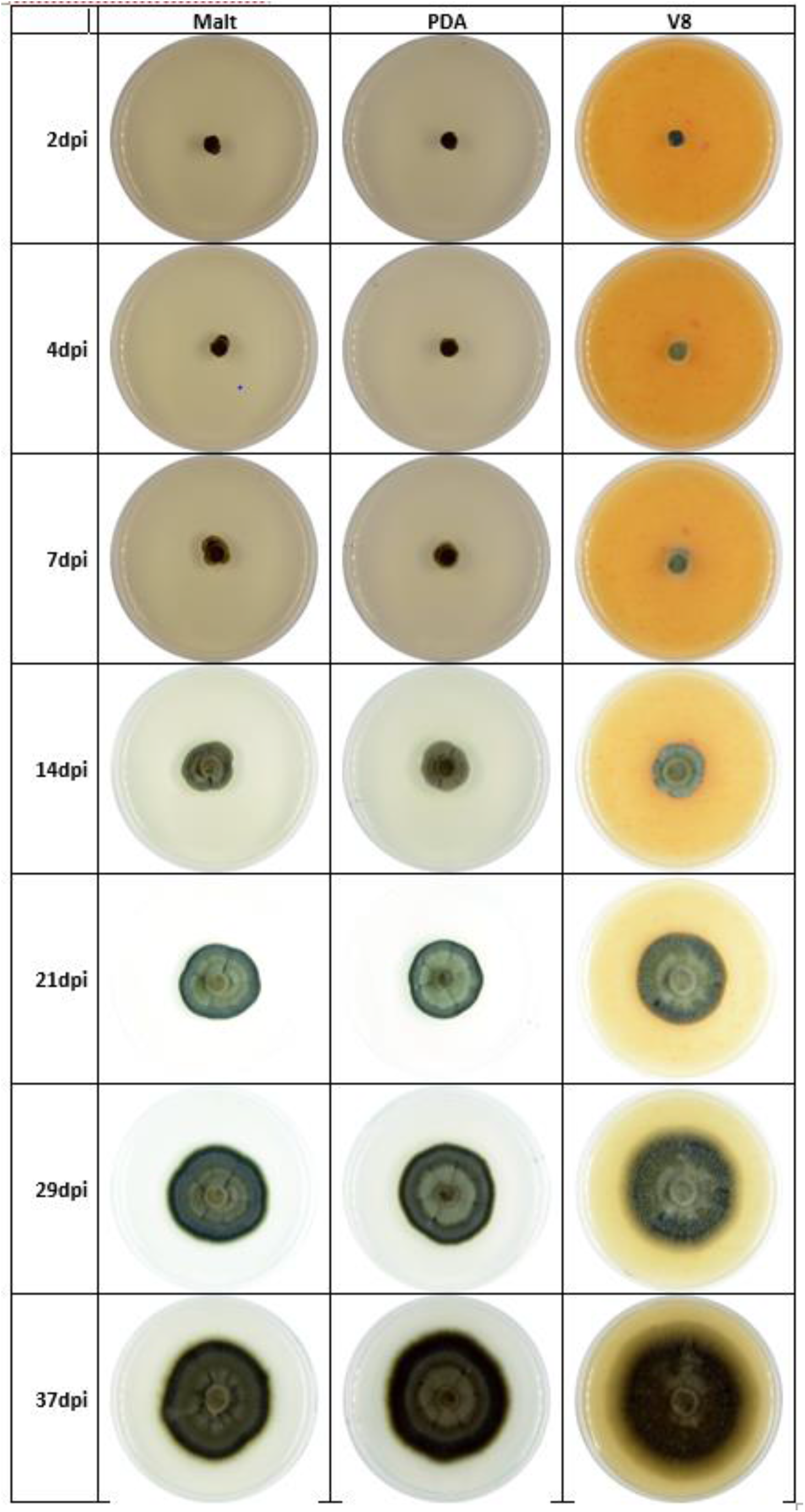
Growth kinetics of the same *Mycosphaerella brassicicola* strain on malt, PDA and V8 agar media.

**Figure 9:**
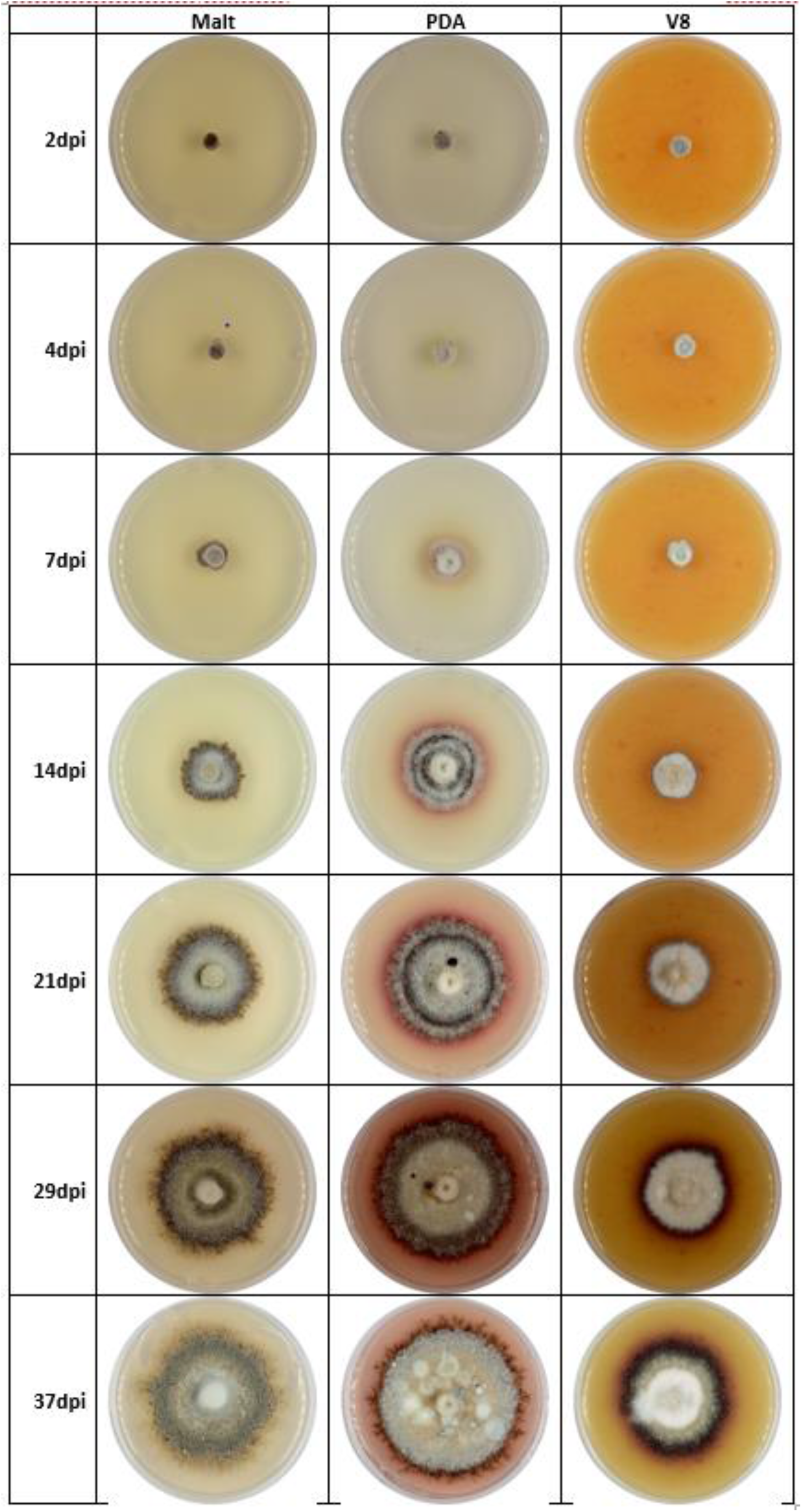
Growth kinetics of the same *Pseudocercosporella capsellae* strain on malt, PDA and V8 agar media.

*P. brassicae* was the slowest (Figure 10). Two weeks were needed to clearly see that the mycelium started growing from the plug, and its colony barely reached 1 cm diameter 37 days after colony transfer.

**Figure 10:**
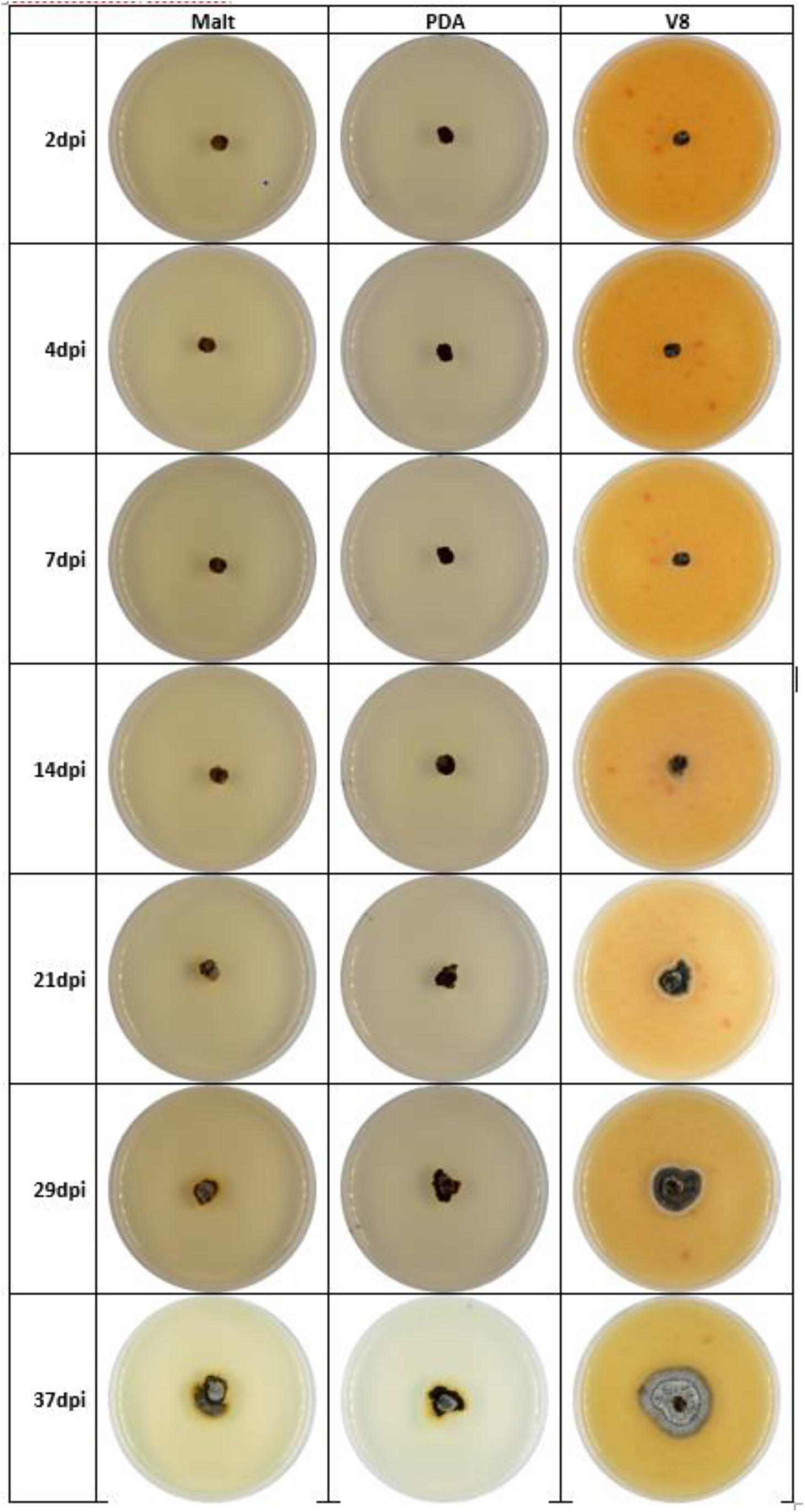
Growth kinetics of the same *Pyrenopeziza brassicae* strain on malt, PDA and V8 agar media.

Noticeably, for the same strain colony growth was be different on the 3 media used. For example, *L. biglobosa* and *L. maculans* grew quicker on V8 and on PDA than on malt (Figure 5, 6); *P. capsellae* grew quicker on PDA than on V8 and on malt (Figure 9). The aspect of the colonies changed over time and depended on the culture medium (Figures 3–10).

### Pigment production

All the *L. biglobosa* strains released a yellow pigment in their second week of growth. This pigment was better seen on malt than on PDA or V8 (Figure 5). This is a criterion useful for the discrimination between *L. biglobosa* and *L. maculans* strains.

Some of the *P. capsellae* strains can release a violet pigment, cercosporin (Gunasinghe et al. 2016). In our culture conditions, cercosporin was produced on PDA but not on malt, and is less visible on V8 (Figure 9).

### Fruiting bodies production

*S. sclerotiorum* was easily recognised by the early production of sclerotia, already well melanised after a week (Figure 3).

It happens that some *L. maculans* and *L. biglobosa* strains spontaneously produce pycnidia. This production can be promoted on V8 medium, exposing the colonies to black (near UV) light.

## Discussion

With the help of the camera stand, taking pictures was quick and easy. This produced homogeneous images, which facilitates their comparison. Nevertheless, a slight colour variation remained, that the post-processing attenuated by white balancing the images.

The speed of colony growth depends on the culture conditions, especially on the temperature (Lovell et al. 2004). This prevent their generalisation, and require considering them with caution. However, the large contrast between species’ growth was evidenced. Similarly, the exact appearance of the colonies changed with the growth medium composition and its physico-chemical characteristics. Specifically, changes in pH or oxydo-reduction level deeply affect the colony aspect, speed of growth and fruiting bodies production (Bousset et al. 2019). However, there again it is the comparison between appearances and the kinetics of change over time that can be retained as discrimination criteria. Having standardised the imaging and having grown the 8 species simultaneously allowed us to propose series of comparative pictures that one can refer to in case of a doubt.

The colony appearance also changes among individuals, as illustrated for *P. brassicae* (Carmody et al. 2020), and for *P. capsellae* (Gunasinghe et al. 2016) or *Leptosphaeria* (Zhang et al. 2014). It will thus always remain necessary to get familiar with the morphological diversity among strains. The availability of standardised pictures may facilitate this learning.

## Acknowledgements

This project benefited from the financial support of INRAE and from the CASDAR ATIPICAL 2019-2022 projet « Actualisation et pérennisation des connaissances et des ressources biologiques et moléculaires sur les maladies foliaires de colza ».

